# Is there an accurate and generalisable way to use soundscapes to monitor biodiversity?

**DOI:** 10.1101/2022.12.19.521085

**Authors:** Sarab S. Sethi, Avery Bick, Robert M. Ewers, Holger Klinck, Vijay Ramesh, Mao-Ning Tuanmu, David A. Coomes

## Abstract

Acoustic monitoring has the potential to deliver biodiversity insight on vast scales. Whilst autonomous recording networks are being deployed across the world, existing analytical techniques struggle with generalisability. This limits the insight that can be derived from audio recordings in regions without ground-truth calibration data. By calculating 128 learned features and 60 soundscape indices of audio recorded during 8,023 avifaunal point counts from diverse ecosystems, we investigated the generalisability of soundscape approaches to biodiversity monitoring. Within each dataset, we found univariate correlations between several acoustic features and avian species richness, but features behaved unpredictably across datasets. Training a machine learning model on compound indices, we could predict species richness within datasets. However, models were uninformative when applied to datasets not used for training. We found that changes in soundscape features were correlated with changes in avian communities across all datasets. However, there were cases where avian communities changed without an associated shift in soundscapes. Our results suggest that there are no common hallmarks of biodiverse soundscapes across ecosystems. Therefore, soundscape monitoring should only be used when high quality ground-truth data exists for the region of interest, and in conjunction with more targeted and accurate in-person ecological surveys. By better understanding how to use interpret data reliably, we hope to unlock the scale at which acoustic monitoring can be used to deliver true impact for land managers and scientists monitoring biodiversity around the world.

**Summary:** Whilst eco-acoustic monitoring has the potential to deliver biodiversity insight on vast scales, existing analytical approaches behave unpredictably across studies. We collated 8,023 audio recordings with paired manual avifaunal point counts to investigate whether soundscapes could be used to monitor biodiversity across diverse ecosystems. We found that neither univariate indices nor machine learning models were predictive of species richness across datasets, but soundscape change was consistently indicative of community change. Our findings indicate that there are no common features of biodiverse soundscapes, and that soundscape monitoring should be used cautiously and in conjunction with more reliable in-person ecological surveys.

## Introduction

Anthropogenic pressures are impacting biodiversity globally^1^. Declines in species richness catch headlines^2^, but changes in community composition can have just as devastating ecological effects^3^. To design evidence-based conservation measures, monitoring biodiversity is essential, and listening to the sounds produced by an ecosystem (acoustic monitoring) holds promise as a scalable and inexpensive way to achieving this^4^.

Many species contribute to an ecosystem’s soundscape, whether it be through producing vocalisations (e.g., birdsong), stridulations (e.g., cricket chirps), or as they interact with the environment (e.g., the buzz of a bee). Streaming^5^ or recording^6^ soundscapes across huge scales is now commonly done^7–9^, yet interpreting the audio to derive biodiversity insight remains a challenge. Automating the identification of stereotyped sounds in audio recordings^10^ can provide species occurrence data on large scales, building a bottom-up picture of biodiversity. However, vocalisation detection algorithms rely on vast amounts of training data^11^, meaning the sonic contributions of all but the most commonly studied species are often ignored. An alternative top-down approach is to use the features of an entire soundscape to infer biodiversity^12^. Entropy-based acoustic indices^12^ or embeddings from ML models^13^ have both been used to predict community richness or ecosystem intactness with some success when calibrated using ground-truth observational data^14,15^. Nonetheless, soundscape features which correlate positively with biodiversity at one site can have an inverse relationship in another^16,17^, and no reliable single metric has been found^18^. The inability of both vocalisation detection and soundscape approaches to generalise has meant acoustic monitoring has only provided ecological insight in already well-studied regions, and the technology’s transformative potential has remained largely unfulfilled.

## Results

### Cross-biome soundscape recordings with associated avian richness

We collated 8,023 short (1-20 minute) soundscape recordings collected concurrently with manually recorded avifaunal community data from four diverse datasets: a temperate forest in Ithaca, USA (N=6,734, one site), a varied tropical rainforest landscape in Sabah, Malaysia (N=977, 14 sites), an agricultural tea landscape in Chiayi, Taiwan (N=165, 16 sites), and a varied montane tropical forest and grassland landscape in the Western Ghats, India (N=147, 91 sites). In India, Malaysia, and Taiwan, avian community data was collected by in-person point counts performed by experts, whereas citizen science checklists from eBird were used for the USA^19^ (Methods). For each recording, we calculated two common types of acoustic features; (i) a 128 dimensional convolutional neural network (CNN) embedding (hereby learned features, LFs)^13,20^ and (ii) 60 analytically derived soundscape indices (SSIs)^21^.

### Directly predicting biodiversity from soundscapes is not generalisable

First, we investigated univariate correlations between soundscape features and avian species richness. We found that many LFs and SSIs correlated with richness (Pearson’s correlation *p<0.05* by permutation test with Bonferroni correction), though the number varied greatly by dataset (**Fig. 1a**). Four learned features and no soundscape indices correlated with avian richness in all four datasets (**Fig. 1b**), whilst most features correlated with richness in either one or two datasets (105 x LFs, 43 x SSIs; **Fig. 1b**). For features that were correlated with avian richness in multiple datasets, the gradient and intercept of fitted lines varied and correlation coefficients were relatively low (e.g., 14^th^ learned feature: *0.048 < r^2^ < 0.286*, ROICover [a measure of spectrogram cover]: *0.002 < r^2^ < 0.099*, **Fig. 1c**).

**Figure 1:**
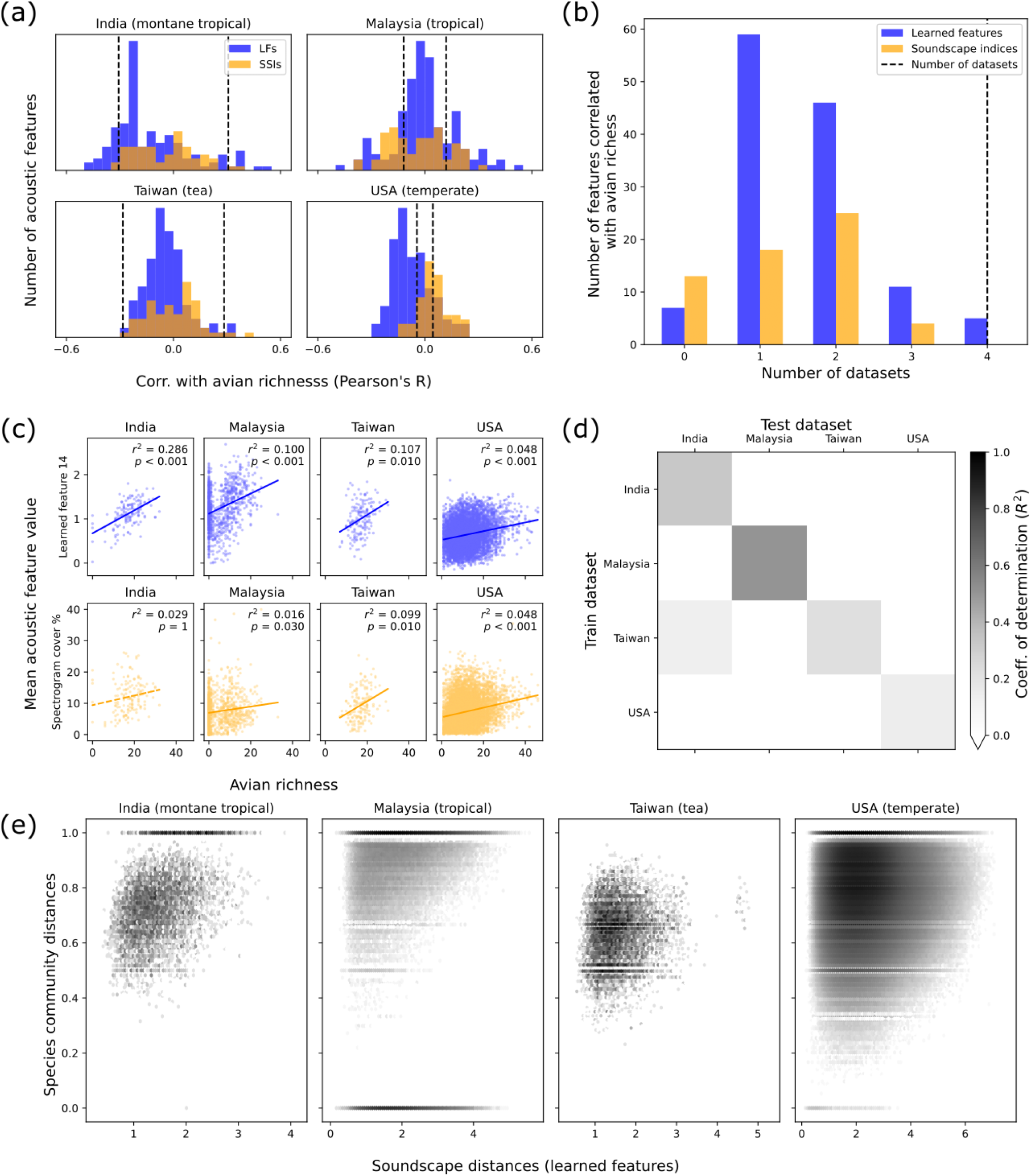
Predictions of avian species richness from acoustic features of soundscapes are not generalisable across datasets, but soundscape change is a reliable indicator of community change. **(a)** Many soundscape indices (SSIs) and learned features (LFs) correlated significantly with avian species richness in each dataset (dashed lines indicate significance thresholds derived from permutation tests). **(b)** However, only four LFs and no SSIs correlated significantly with richness in all datasets, with most only correlating with richness in one or two datasets (105 LFs, 43 SSIs). **(c)** For spectrogram cover and LF 14 (both of which correlated with richness in many datasets) the gradient and intercept of the correlations varied significantly between datasets, and correlation coefficients were relatively low, limiting transferability. **(d)** Predicting avian richness with a machine learning model trained on LFs was moderately successful when training and test datasets were the same. However, models were unable to generalise when predicting richness in datasets other than the one used for training. **(e)** For each dataset we found that pairwise distances between the acoustic features of soundscapes correlated with Jaccard distances between avian communities present. With Jaccard distances, 0 indicates identical communities and 1 indicates no community overlap.

By training a machine learning model on the full dimensional learned feature vectors, we were able to predict species richness within datasets with relative success (coefficient of determination *R^2^ = 0.32, 0.50, 0.22, 0.14* for India, Malaysia, Taiwan, USA, respectively; **Fig. 1d**). However, the models were unable to predict richness when evaluated on datasets that they were not trained upon (**Fig. 1d**). Our results indicate that even within a single dataset – but especially when looking across datasets – soundscapes with similar levels of avian diversity do not share similar acoustic features.

### Soundscape change consistently correlates with avian community change

Rather than attempting to predict species richness directly, we investigated the relationship between pairwise Euclidean distances between the mean acoustic features of each audio recording (soundscape change), and pairwise Jaccard distances between their associated avian species communities (community change). We found strong correlations between soundscape change and community change within each dataset (Spearman correlation Mantel test, *p ≤ 0.001* for all; **Fig. 1e**). There were more examples where communities changed but soundscapes didn’t than the converse, indicated by clustering of points in the upper left of **Fig. 1e**. No significant correlations were found between soundscape change and change in species richness in any of the four datasets (SI Fig. S2). For each of the 14 sites in Malaysia – the only dataset with sufficient replicates at individual sites to test for this relationship – there was a significant correlation between soundscape change and community change (*p ≤ 0.03*). The decision to use learned features (**Fig. 1e**) or soundscape indices (SI Fig. S3) to track soundscape change did not alter our findings.

## Discussion

In all four datasets, we found that certain acoustic features correlated with avian richness. However, features did not behave consistently across datasets, and most were only correlated with richness in one or two datasets (explaining inconsistencies seen in the literature^18^, **Figs. 1a, 1b, 1c**). Without access to ground-truth calibration data, ascertaining which features were most suitable for each dataset would be impossible, limiting the transferability of this approach. We then found that a machine learning model trained on compound indices was able to produce within-dataset predictions of species richness with some success (**Fig. 1d**). But, again, generalisability was not achieved as models did not produce informative estimates when applied to datasets they were not trained upon. The diversity of biophony, anthropophony, and geophony in soundscapes across the diverse datasets would, perhaps, make these results unsurprising. Nonetheless, they stress that however well an acoustic feature or ML model may perform in certain scenarios, without access to high quality ground truth data from the exact location being studied, soundscape methods should not be used to generate predictions of species richness.

In contrast to the unpredictable behaviour of the models producing estimates of avian richness, we found that soundscape change correlated with community change in all datasets (**Fig. 1e**). However, this approach was also limited, as there were many examples where similar soundscapes were associated with very different avian communities. One reason could be that non-biotic sounds (e.g., motors) or non-avian species (e.g., insects, amphibians) were larger contributors to soundscapes than birds^14,22^. Our ability to predict avian richness was therefore more likely to have been based on latent variables measuring habitat suitability (e.g., vocalising prey, or nearby water sources) rather than being driven the vocal contributions of birds directly.

Despite having access to vast amounts of ground-truth data, even within datasets, soundscape predictions of avian richness and community change struggled with accuracy. Rather than relying on these predictions in isolation, a more reliable approach might be to use soundscapes to collect coarse but scalable data, which can be used to direct more targeted ecological studies. This approach could result in more efficient use of limited expert resources to ensure interventions are deployed in a timely and focussed manner. Indeed, due to issues surrounding interpretability and accountability, we will always likely require expert verification of autonomous monitoring outputs before policy and management practices are modified^23^. Proceeding with more realistic expectations around how soundscapes can best contribute large-scale biodiversity monitoring efforts will be essential to maximising the transformative potential of the technology.

## Acknowledgements

We would like to thank staff at the Stability of Altered Forest Ecosystems (SAFE) project, the Cornell Lab of Ornithology, and Ecoacoustics and Spatial Ecology Lab at Academia Sinica for their support in data collection and pre-processing. Specific thanks go to Jani Sleutel, Adi Shabrani, Nursyamin Zulkifli, Ray Mack, Ben Thomas, Akshay Anand, Chandrasekar Das, Amrutha Rajan, Jia-Yun Lee, Shih-Hao Liu.

This project was supported by funding from the Herchel Smith Fund (SSS), World Wildlife Fund (Malaysia data), Sime Darby Foundation (Malaysia data), National Science and Technology Council (NSTC 111-2321-B-002-019 and 111-2927-I-001-513; Taiwan data), Biodiversity Research Center at Academia Sinica (Taiwan data). Feature computations for the USA dataset were performed on resources provided by Sigma2 - the National Infrastructure for High Performance Computing and Data Storage in Norway. Data was collected from Malaysia under an SaBC permit granted to SSS (JKM/MBS.1000-2/2 JLD.8 (63)).

## Data availability

Processed acoustic features and avian community data for all point counts is published at https://doi.org/10.5281/zenodo.7410357. Code with instructions to reproduce analyses, results, and figures can be found at https://doi.org/10.5281/zenodo.7458688.

## Methods

### Avifaunal point counts

#### India

Data was collected from 91 sites in the Western Ghats between March 2020 and May 2021. Sites were a mixture of montane wet evergreen and semi-evergreen forests (29), montane grasslands (8), moist deciduous forests (7), timber plantations (17), tea plantations (20), agricultural land (8), and settlements (2).

147 15-minute point counts were conducted between 6-10am at each site (mean 1.6 per site). A variable-distance point count approach was followed, and all bird species heard, seen, and those that flew over (primarily raptor species) were noted. All point counts were carried out between 6am and 10am (timing of high avian activity) at each location.

Single channel audio was recorded during each point count using an Audiomoth device^6^. Recordings were saved in WAV format at a sampling rate of 48 kHz.

#### Malaysia

Data was collected from a varied tropical landscape at the Stability of Altered Forest Ecosystems (SAFE) project^24^ in Sabah between March 2018 and February 2020. The 14 sites spanned a degradation gradient: two in protected old growth forest, two in a protected riparian reserve, six in selectively logged forest (logging events in 1970s and early 2000s), two in salvage logged forest (last logged in early 2010s), and two in oil palm plantations. Sites had a mean separation distance of 7.6km, with the closest two being 583m apart.

977 20-minute avifaunal point counts were performed across 24 hours of the day at each site (59-80 per site). During point counts all visual or aural encounters of avifaunal species within a 10m radius of the sampling site were recorded. Species identifications and names were validated using the Global Biodiversity Information Facility^25^ (GBIF). A total of 216 avifaunal species were recorded across all point counts.

Single channel audio was recorded during each point count with a Tascam DR-05 recorder mounted to a tripod at chest height (nominal input level -20 dBV, range 20Hz-22kHz). Recordings were saved in WAV format at a sampling rate of 44.1k kHz.

#### Taiwan

Data was collected from 16 tea plantations in the Alishan tea district, located in Chiayi County of Taiwan, between January and November 2022. The tea plantations spanned an elevation gradient from 816-1464 m and were surrounded by secondary broadleaf forests or coniferous plantations.

Stereo audio was recorded at a sampling rate of 44.1kHz for one of every fifteen minutes at each site throughout the sampling period. Wildlife Acoustics Song Meter 4 devices were used (mounted to a tree truck or tripod at chest height) and data was saved in WAV format.

176 10-minute avifaunal point counts were conducted in total, on average once per site per month. Every survey was conducted within 3 hours after sunrise. During point counts, the species of every bird individual visually or aurally detected was recorded and its horizontal distance from the observer was estimated as either 0-25m, 25-50m, 50-100m, >100m or flying over. Our process for matching audio recordings to point counts is provided later in the Methods.

#### USA

Data was collected from a single site in a temperate forest at Sapsucker Woods, Ithaca from January 2016 to December 2021.

Audio was recorded continuously using a Gras-41AC precision microphone mounted at chest height and digitised with a Barix Instreamer ADC at a sampling rate of 48 kHz. Files were saved in 15-minute chunks and in WAV format.

In the absence of standardised point counts across such a long duration, we used eBird checklists to determine avian communities (only possible since Sapsucker Woods is a major hotspot for eBird data). Data was filtered to only keep checklists which were complete, of the “travelling” or “stationary” types, located within 200m of the recording location, and shorter than 30 minutes. We combined occurrence data from all eBird checklists that started during each audio recording to create one pseudo point count per audio file. Since we only use occurrence data, double counting of the same species was not an issue. This resulted in a total of 6,734 point counts.

### Avian communities

For each point count, we used the field observations to derive community occurrence data (i.e., presence or absence) since not all datasets had reliable abundance information. To calculate avian richness, we calculated the unique number of species encountered within each point count (α-diversity).

### Acoustic features

We used two approaches to calculate acoustic features for each recording. The first was using a suite of 60 soundscape indices (SSIs), using the *sciki-maad* library^21^. Specifically, we appended together the feature vectors from the functions *maad.features.all_temporal_alpha_indices* and *maad.features.all_spectral_alpha_indices*.

The second approach was to use a learned feature embedding (LF) from the VGGish convolutional neural network (CNN) model^20^. VGGish is a pre-trained, general purpose sound classification model that was trained on the AudioSet database^26^. First audio was downsampled to 16kHz then converted into a log-scaled mel-frequency spectrogram (window size = 25ms, window hop = 10ms, periodic Hann window). Frequencies were mapped to 64 mel-frequency bins between 125-7500Hz, and magnitudes were offset before taking the logarithm. Data was inputted to the CNN in spectrograms of dimension 96×64, producing one 128-dimensional acoustic feature vector for each 0.96s of audio.

For all acoustic features (LFs and SSIs) mean feature vectors were calculated for each point count. For Malaysia, India, and USA, recordings had a 1:1 mapping with point counts, so features were averaged across each audio recording. For Taiwan, where audio data was sparser, features were averaged across all audio recordings that begun within a one-hour window centred around the point count’s start time, resulting in an average of four 1-minute audio recordings per point count.

### Univariate correlations with avian richness

Univariate Pearson correlations were calculated between each of the 60 SSIs and 128 LFs with avian richness across all point counts within datasets. Significance thresholds were determined by calculating 100 null correlation coefficients between shuffled features and avian richness. Null correlation coefficients were aggregated for each dataset across all 60 SSIs and 128 LFs (total null values = 18800). Accounting for a Bonferroni multiple hypothesis correction, the threshold for significance at *p=0.05* was taken as value 18784 in an array of sorted absolute null correlation coefficients in ascending order. For each dataset, all real correlations between acoustic features and avian richness with an absolute correlation coefficient greater than this threshold were determined to be significant. Lines of best fit in Fig. 1c were determined by fitting a 1^st^ order polynomial to the avian richness (x-axis) and acoustic feature (y-axis) data.

### Machine learning predictions of avian richness

We used a random forest regression model to make predictions of avian richness using the full dimensional acoustic feature vectors. The *scikit-learn* implementation *RandomForestRegressor* was used with a fixed random seed and other default parameters unchanged. We randomly selected 70% of point counts in each dataset for training and held back 30% for testing. We measured how well predicted richness matched true richness using the coefficient of determination (*R^2^*). Both within and across datasets (i.e., for all results in Fig. 1d, SI Fig. S1), models were always trained on the training data and scores were evaluated on the test data.

### Investigating link between soundscape change and community change

To measure community change between two point counts we used the Jaccard distance metric, where 0 indicates no change in community and 1 indicates no shared species between communities. To measure soundscape change we calculated the Euclidean distance between the mean acoustic feature vectors for each point count. Performing these two operations between all point counts within each dataset, we derived two matrices representing pairwise community change and pairwise soundscape change. Since community change was bounded between 0 and 1, a Pearson correlation was not appropriate. Therefore, the Spearman correlation between these two matrices was measured using a one-tailed Mantel test, with p-values derived empirically (using the *skbio* implementation *skbio.stats.distance.mantel*).

## Supplementary Information

**Figure S1:**
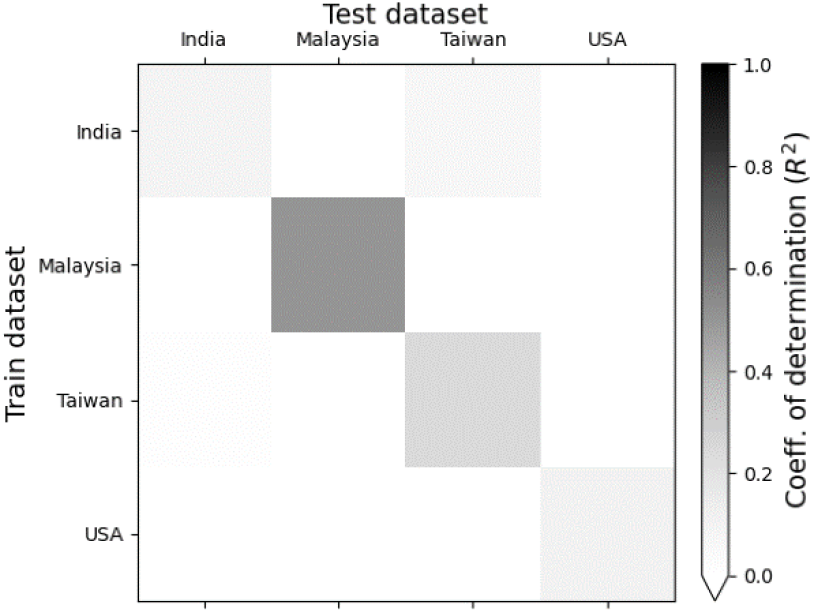
When training and test datasets were the same, predictions of species richness from a machine learning model (Random Forest Regressor) trained on soundscape indices (rather than learned features in Fig. 1c) were of varying levels of accuracy (R^2^=0.09, 0.50, 0.24, 0.10 for India, Malaysia, Taiwan, USA, respectively). In all cases, predictive models did not generalise to produce informative estimates of species richness when applied to datasets they were not trained on.

**Figure S2:**
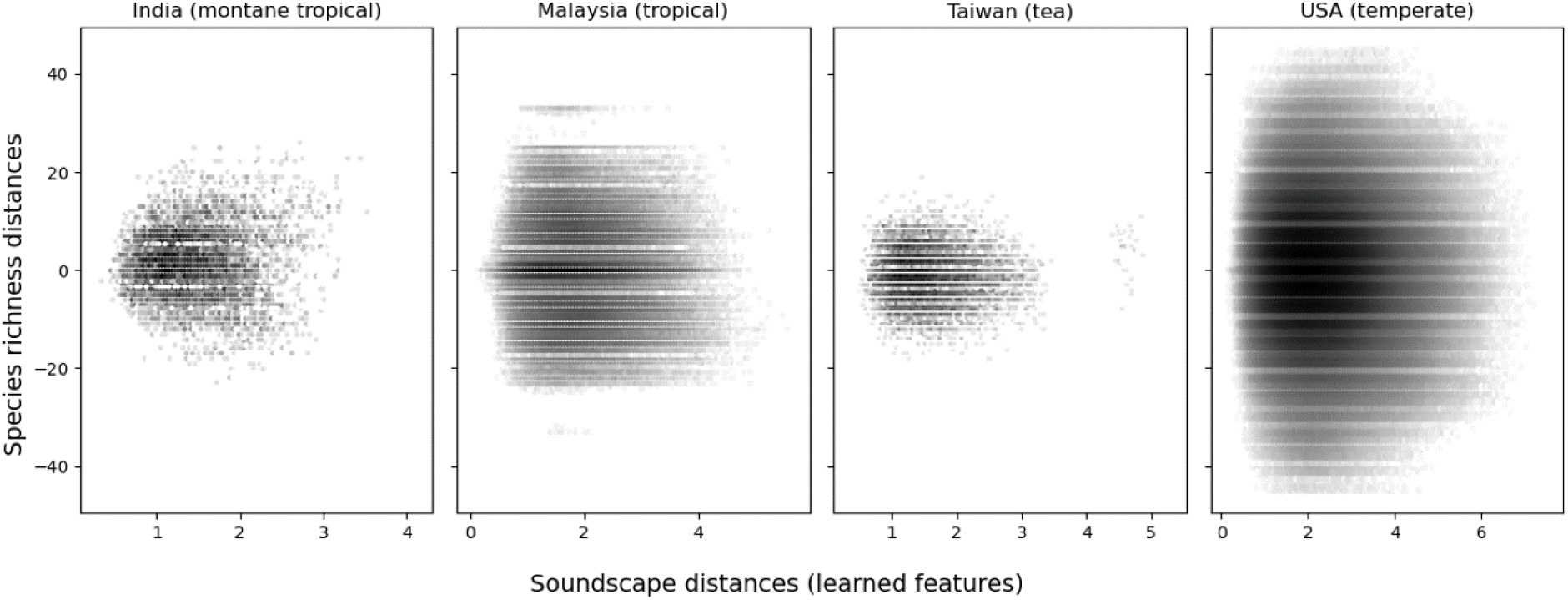
No significant correlations were found between change in soundscape features and change in species richness in any of the four datasets (Spearman’s correlation Mantel test, *p > 0.05* in all cases).

**Figure S3:**
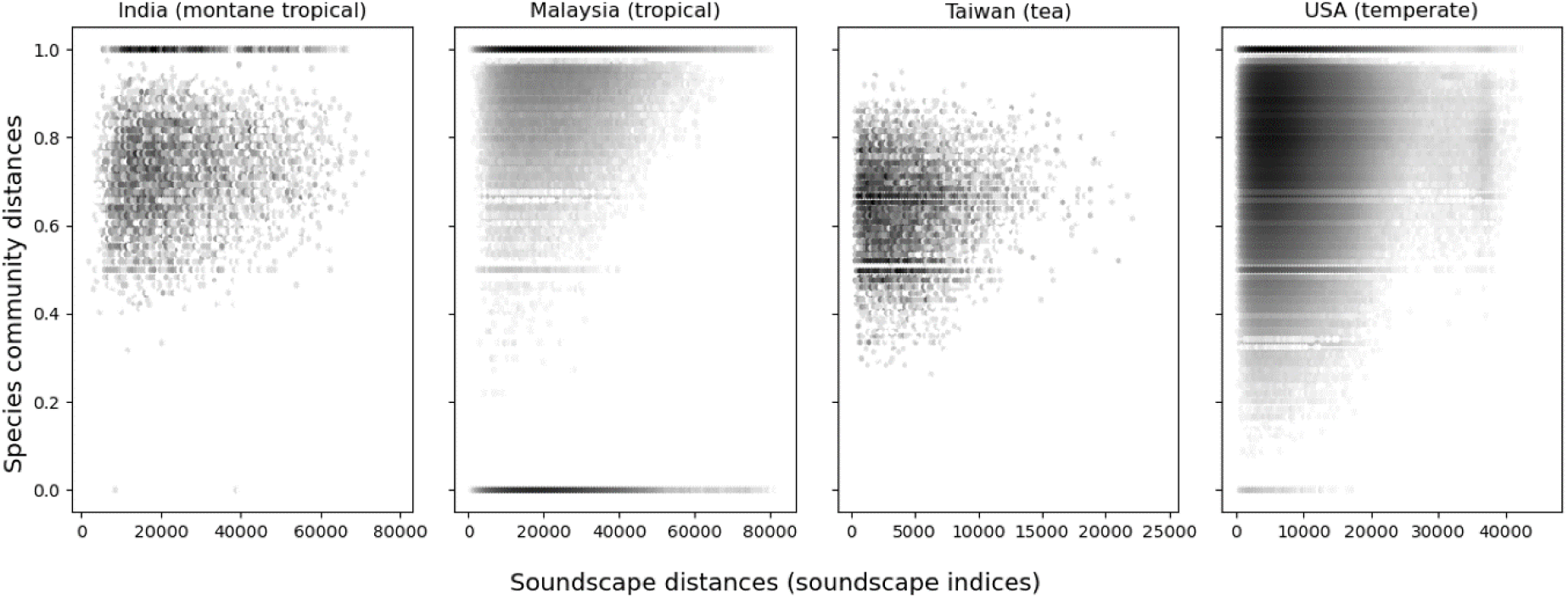
Even when using soundscape indices rather than learned acoustic features, change in soundscape features was correlated with change in avian community across all datasets (Spearman’s correlation Mantel test, *p = 0.002, 0.001, 0.023, 0.001* for India, Malaysia, Taiwan, USA).

## References

1. Pereira, H. M., Navarro, L. M. & Martins, I. S. Global Biodiversity Change: The Bad, the Good, and the Unknown. Annu. Rev. Environ. Resour. 37, 25–50 (2012).

2. Butchart, S. H. M. et al. Global biodiversity: Indicators of recent declines. Science 328, 1164–1168 (2010).

3. Pereira, H. M. & Cooper, H. D. Towards the global monitoring of biodiversity change. Trends Ecol. Evol. 21, 123–129 (2006).

4. Gibb, R., Browning, E., Glover-Kapfer, P. & Jones, K. E. Emerging opportunities and challenges for passive acoustics in ecological assessment and monitoring. Methods Ecol. Evol. 10, 169–185 (2019).

5. Sethi, S. S., Ewers, R. M., Jones, N. S., Orme, C. D. L. & Picinali, L. Robust, real-time and autonomous monitoring of ecosystems with an open, low-cost, networked device. Methods Ecol. Evol. 9, 2383–2387 (2018).

6. Hill, A. P. et al. AudioMoth: Evaluation of a smart open acoustic device for monitoring biodiversity and the environment. Methods Ecol. Evol. 9, 1199–1211 (2018).

7. Roe, P. et al. The Australian Acoustic Observatory. Methods Ecol. Evol. 12, 1802–1808 (2021).

8. Sethi, S. S. et al. SAFE Acoustics: An open-source, real-time eco-acoustic monitoring network in the tropical rainforests of Borneo. Methods Ecol. Evol. 11, 1182–1185 (2020).

9. Wood, C. M., Kahl, S., Chaon, P., Peery, M. Z. & Klinck, H. Survey coverage, recording duration and community composition affect observed species richness in passive acoustic surveys. Methods Ecol. Evol. 12, 885–896 (2021).

10. Stowell, D. Computational bioacoustics with deep learning: a review and roadmap. PeerJ 10, e13152 (2022).

11. Kahl, S., Wood, C. M., Eibl, M. & Klinck, H. BirdNET: A deep learning solution for avian diversity monitoring. Ecol. Inform. 61, 101236 (2021).

12. Sueur, J., Pavoine, S., Hamerlynck, O. & Duvail, S. Rapid acoustic survey for biodiversity appraisal. PLOS ONE 3, e4065 (2008).

13. Sethi, S. S. et al. Characterizing soundscapes across diverse ecosystems using a universal acoustic feature set. Proc. Natl. Acad. Sci. 117, 17049–17055 (2020).

14. Sethi, S. S. et al. Soundscapes predict species occurrence in tropical forests. Oikos 2022, e08525 (2022).

15. Sueur, J., Farina, A., Gasc, A., Pieretti, N. & Pavoine, S. Acoustic Indices for Biodiversity Assessment and Landscape Investigation. (2014) doi:info:doi/10.3813/AAA.918757.

16. Mammides, C., Goodale, E., Dayananda, S. K., Kang, L. & Chen, J. Do acoustic indices correlate with bird diversity? Insights from two biodiverse regions in Yunnan Province, south China. Ecol. Indic. 82, 470–477 (2017).

17. Bohnenstiehl, D., Lyon, R., Caretti, O., Ricci, S. & Eggleston, D. Investigating the utility of ecoacoustic metrics in marine soundscapes. J. Ecoacoustics 2, R1156L (2018).

18. Alcocer, I., Lima, H., Sugai, L. S. M. & Llusia, D. Acoustic indices as proxies for biodiversity: a meta-analysis. Biol. Rev. 97, 2209–2236 (2022).

19. Sullivan, B. L. et al. The eBird enterprise: An integrated approach to development and application of citizen science. Biol. Conserv. 169, 31–40 (2014).

20. Hershey, S. et al. CNN architectures for large-scale audio classification. in 2017 IEEE International Conference on Acoustics, Speech and Signal Processing (ICASSP) 131–135 (2017). doi:10.1109/ICASSP.2017.7952132.

21. Ulloa, J. S., Haupert, S., Latorre, J. F., Aubin, T. & Sueur, J. scikit-maad: An open-source and modular toolbox for quantitative soundscape analysis in Python. Methods Ecol. Evol. 12, 2334–2340 (2021).

22. Ferreira, L. et al. What do insects, anurans, birds, and mammals have to say about soundscape indices in a tropical savanna. J. Ecoacoustics 2, PVH6YZ (2018).

23. Smith, J. W. & Pijanowski, B. C. Human and policy dimensions of soundscape ecology. Glob. Environ. Change 28, 63–74 (2014).

24. Ewers, R. M. et al. A large-scale forest fragmentation experiment: The Stability of Altered Forest Ecosystems Project. Philos. Trans. R. Soc. Lond. B Biol. Sci. 366, 3292–3302 (2011).

25. GBIF Secretariat. GBIF backbone taxonomy. Checkl. Dataset Accessed March 2020 (2020).

26. Gemmeke, J. F. et al. Audio Set: An ontology and human-labeled dataset for audio events. in 2017 IEEE International Conference on Acoustics, Speech and Signal Processing (ICASSP) 776–780 (2017). doi:10.1109/ICASSP.2017.7952261.

